# Combined Treatment with Dopamine Receptor Antagonists and Radiation Creates a Metabolic Vulnerability in Mouse Models of Glioblastoma

**DOI:** 10.1101/2020.01.13.905380

**Authors:** Mohammad Saki, Kruttika Bhat, Fei Cheng, Ling He, Le Zhang, Angeliki Ioannidis, David Nathanson, Jonathan Tsang, Phioanh Leia Nghiemphu, Timothy F. Cloughesy, Linda M. Liau, Harley I. Kornblum, Frank Pajonk

## Abstract

**Background:** Glioblastoma is the deadliest brain tumor in adults and the standard-of-care consists of surgery followed by radiation and treatment with temozolomide. Overall survival times for patients suffering from glioblastoma are unacceptably low indicating an unmet need for novel treatment options.

**Methods:** Using patient-derived glioblastoma lines and mouse models of glioblastoma we test the effect of radiation and the dopamine receptor antagonist on glioblastoma self-renewal *in vitro* and survival *in vivo.* A possible resistance mechanism is investigated using RNA-Sequencing.

**Results:** Treatment of glioma cells with the dopamine receptor antagonist quetiapine reduced glioma cell self-renewal *in vitro* and combined treatment of mice with quetiapine and radiation prolonged the survival of glioma-bearing animals. The combined treatment induced the expression of genes involved in cholesterol biosynthesis. This rendered the tumors vulnerable to simultaneous treatment with atorvastatin and further significantly prolonged the survival of the animals.

**Conclusions:** Our results indicate high efficacy of a triple combination of quetiapine, atorvastatin and radiation against glioblastoma without increasing the toxicity of radiation. With both drugs readily available for clinical use our study could be rapidly translated into a clinical trial.

## Introduction

Despite decades of drug development and technical improvement in radiotherapy, glioblastoma is still the deadliest brain cancer in adults with almost all patients ultimately dying from the disease (1). Attempts to add chemotherapeutic drugs that serve as radiosensitizers have largely failed due to either lack of blood-brain-barrier (BBB) penetration or lack of a proper therapeutic window with temozolomide being the only radiosensitizer that has so far been included into the standard-of-care (2). The realization that glioblastoma cell populations in a tumor are heterogenous with respect to their clonal origin but also their ability to initiate tumors, techniques to identify glioma-initiating cells prospectively (3, 4), and the inherent radioresistance of glioma-initiating cells (5) have sparked research aiming to target glioma-initiating cells more specifically (6).

In a recent high-throughput screen of 83,000 compounds we identified the first-generation dopamine receptor antagonists trifluoperazine as an FDA-approved drug with known BBB penetration that interferes with the self-renewal of glioma cells alone and in combination with radiation (7). Considering the unfavorable side effect profile of trifluoperazine we have extended our studies to include second-generation dopamine receptor antagonists with milder side effects.

We report here that Quetiapine (QTP), like trifluoperazine (TFP), when combined with radiation reduced the self-renewal of patient-derived glioblastoma specimens *in vitro* and prolonged survival in mouse models of glioblastoma. We describe, that the combined treatment induced the expression of genes involved in cholesterol biosynthesis, thus creating a metabolic vulnerability that could be exploited by the use of the statin atorvastatin leading to significantly improved survival.

## Material and Methods

### Cell culture

Primary human glioma cell lines were established at UCLA as described (Hemmati *et al*., PNAS 2003 (3); Characteristics of specific gliomasphere lines can be found in Laks *et al*., Neuro-Oncology 2016 (8)). The GL261 murine glioma cell line was from obtained from Charles River Laboratories, Inc., Frederick, MD. GL261 cells were cultured in log-growth phase in DMEM (Invitrogen, Carlsbad, CA) supplemented with 10% fetal bovine serum, penicillin and streptomycin. Primary glioblastoma cells were propagated as gliomaspheres in serum-free conditions in ultra-low adhesion plates in DMEM/F12, supplemented with B27, EGF, bFGF and heparin as described previously (9). All cells were grown in a humidified atmosphere at 37°C with 5% CO_2_. The unique identity of all patient-derived specimen was confirmed by DNA fingerprinting (Laragen, Culver City, CA). All lines were routinely tested for mycoplasma infection (MycoAlert, Lonza).

### Animals

6–8-week-old C57BL/6 mice, or NOD-*scid* IL2Rgamma^null^ (NSG) originally obtained from The Jackson Laboratories (Bar Harbor, ME) were re-derived, bred and maintained in a pathogen-free environment in the American Association of Laboratory Animal Care-accredited Animal Facilities of Department of Radiation Oncology, University of California (Los Angeles, CA) in accordance to all local and national guidelines for the care of animals. Weights of the animals were recorded every day. 2×10^5^ GL261-Luc or 3×10^5^ HK-374-Luc cells were implanted into the right striatum of the brains of mice using a stereotactic frame (Kopf Instruments, Tujunga, CA) and a nano-injector pump (Stoelting, Wood Dale, IL). Injection coordinates were 0.5mm anterior and 2.25mm lateral to the bregma. The needle was placed at a depth of 3.5 mm from the surface of the brain and retracted 0.5 mm for injection. Tumors were grown for 3 (HK-374), 7 (GL261) days after which successful grafting was confirmed by bioluminescence imaging.

### Drug treatment

After confirming tumor grafting via bioluminescence imaging, mice implanted with the HK-374 or GL261 specimen were injected subcutaneously (quetiapine) or intraperitoneally (Atorvastatin) on a 5-days on / 2-days off schedule with Quetiapine, combined Quetiapine and Atorvastatin, or saline until they reached euthanasia endpoints. Quetiapine was dissolved in acidified PBS (0.4% Glacial acetic acid) at concentration of 5 mg/ml. Atorvastatin was dissolved in corn oil containing 2.5% DMSO at concentration of 5 mg/ml. All animals were treated with 30 mg/kg quetiapine. Atorvastatin was applied by i.p. injection at 30 mg/kg.

### Mass spectrometry

#### Sample Preparation

Whole blood from mice was centrifuged to isolate plasma. Quetiapine was isolated by liquid-liquid extraction from plasma: 50 *µ*l plasma was added to 2 *µ*l internal standard and 100 *µ*l acetonitrile. Mouse brain tissue was washed with 2 mL cold PBS and homogenized using a tissue homogenizer with fresh 2 mL cold PBS. Quetiapine was then isolated and reconstituted in a similar manner by liquid-liquid extraction: 100 *µ*l brain homogenate was added to 2 *µ*l internal standard and 200 *µ*l acetonitrile. After vortex mixing, the samples were centrifuged. The supernatant was removed and evaporated by a rotary evaporator and reconstituted in 100 *µ*l 50:50 water:acetonitrile.

#### Quetiapine Detection

Chromatographic separations were performed on a 100 × 2.1 mm Phenomenex Kinetex C18 column (Kinetex) using the 1290 Infinity LC system (Agilent). The mobile phase was composed of solvent A: 0.1% formic acid in Milli-Q water, and B: 0.1% formic acid in acetonitrile. Analytes were eluted with a gradient of 5% B (0-4 min), 5-99% B (4-32 min), 99% B (32-36 min), and then returned to 5% B for 12 min to re-equilibrate between injections. Injections of 20 *µ*l into the chromatographic system were used with a solvent flow rate of 0.10 ml/min.

Mass spectrometry was performed on the 6460 triple quadrupole LC/MS system (Agilent). Ionization was achieved by using electrospray in the positive mode and data acquisition was made in multiple reactions monitoring (MRM) mode. Two MRM transitions were used for quetiapine: m/z 384.2→ 253.1 and 284.2→ 279.1 with a fragmentor voltage of 140V, and collision energy of 16 and 20 eV, respectively. The analyte signal was normalized to the internal standard and concentrations were determined by comparison to the calibration curve (0.5, 5, 50, 250, 500, 2000 nM). Quetiapine brain concentrations were adjusted by 1.4% of the mouse brain weight for the residual blood in the brain vasculature as described by Dai et al. (10).

### Irradiation

Cells were irradiated at room temperature using an experimental X-ray irradiator (Gulmay Medical Inc. Atlanta, GA) at a dose rate of 5.519 Gy/min for the time required to apply a prescribed dose. The X-ray beam was operated at 300 kV and hardened using a 4mm Be, a 3mm Al, and a 1.5mm Cu filter and calibrated using NIST-traceable dosimetry. Corresponding controls were sham irradiated.

Mice were irradiated using an image-guided small animal irradiator (X-RAD SmART, Precision X-Ray, North Branford, CT) with an integrated cone beam CT (60 kVp, 1 mA) and a bioluminescence-imaging unit. The X-ray beam was operated at 225KV and calibrated with a micro-ionization chamber using NIST-traceable dosimetry. Cone beam CT images were acquired with a 2mm Al filter. For delivery of the radiation treatment the beam was hardened using a 0.3mm Cu filter.

During the entire procedure the interior of the irradiator cabinet was maintained at 35° C to prevent hypothermia of the animals. Anesthesia of the animals was initiated in an induction chamber. Once deeply anesthetized, the animals were then immobilized using custom 3D-printed mouse holder (MakerBot Replicator+, PLA filament) with ear pins and teeth bar, which slides onto the irradiator couch. Cone beam CT images were acquired for each individual animal. Individual treatment plans were calculated for each animal using the SmART-Plan treatment planning software (Precision X-Ray). Radiation treatment was applied using a square 1×1cm collimator from a lateral field. For the assessment of the effect of quetiapine and quetiapine plus Atorvastatin in combination with irradiation *in vivo*, mice were treated with corresponding drugs 1 hour prior to irradiation. Animals received a single dose of 10 Gy on day 3 or day 7 after tumor implantation.

### *In-vitro* sphere formation assay

For the assessment of self-renewal *in vitro*, cells were irradiated with 0, 2, 4, 6 or 8 Gy and seeded under serum-free conditions into untreated plates in DMEM/F12 media, supplemented with 10 ml / 500 mL of B27 (Invitrogen), 0.145 U/ml recombinant insulin (Eli Lilly, Indiana), 0.68 U/mL heparin (Fresenius Kabi, Illinois), 20 ng/ml fibroblast growth factor 2 (bFGF, Sigma) and 20 ng/ml epidermal growth factor (EGF, Sigma). The number of spheres formed at each dose point was normalized against the non-irradiated control. The resulting data points were fitted using a linear-quadratic model.

### Quantitative Reverse Transcription-PCR

Total RNA was isolated using TRIZOL Reagent (Invitrogen). cDNA synthesis was carried out using the SuperScript Reverse Transcription IV (Invitrogen). Quantitative PCR was performed in the QuantStudio™ 3 Real-Time PCR System (appliedbiosystems) using the PowerUp™ SYBR™ Green Master Mix (Applied Biosystems, Carlsbad, CA, USA). *C*_t_ for each gene was determined after normalization to IPO8, TBP, PPIA and ΔΔ*C*_t_ was calculated relative to the designated reference sample. Gene expression values were then set equal to 2^−ΔΔCt^ as described by the manufacturer of the kit (Applied Biosystems). All PCR primers were synthesized by Invitrogen and designed for the human sequences of cholesterol biosynthesis pathway genes: DHCR7, HMGCR, HMGCS1, MVD, SQLE, SREBF2, etc. and PPIA, TBP and IPO8 as housekeeping genes (for primer sequences see **Supplementary Table1**).

### RNASeq

48 hours after 4 Gy irradiation or sham irradiation, RNA was extracted from HK-374 cells using Trizol. RNASeq analysis was performed by Novogene (Chula Vista, CA). Quality and integrity of total RNA was controlled on Agilent Technologies 2100 Bioanalyzer (Agilent Technologies; Waldbronn, Germany). The RNA sequencing library was generated using NEBNext® Ultra RNA Library Prep Kit (New England Biolabs) according to manufacturer’s protocols. The library concentration was quantified using a Qubit 3.0 fluorometer (Life Technologies), and then diluted to 1 ng/*µ*l before checking insert size on an Agilent Technologies 2100 Bioanalyzer (Agilent Technologies; Waldbronn, Germany) and quantifying to greater accuracy by quantitative Q-PCR (library molarity >2 nM). The library was sequenced on the Illumina NovaSeq6000 platform.

Downstream analysis was performed using a combination of programs including STAR, HTseq, and Cufflink. Alignments were parsed using the program Tophat and differential expressions were determined through DESeq2. Reference genome and gene model annotation files were downloaded from genome website browser (NCBI/UCSC/Ensembl) directly. Indexes of the reference genome were built using STAR and paired-end clean reads were aligned to the reference genome, using STAR (v2.5). HTSeq v0.6.1 was used to count the read numbers mapped of each gene. The FPKM of each gene was calculated based on the length of the gene and reads count mapped to this gene.

Differential expression analysis between irradiated and control samples (three biological replicates per condition) was performed using the DESeq2 R package (2_1.6.3). The resulting *p*-values were adjusted using the Benjamini and Hochberg’s approach for controlling the False Discovery Rate (FDR). Genes with an adjusted *p*-value of <0.05 found by DESeq2 were assigned as differentially expressed. To identify the correlations between differentially expressed genes, we generated heatmaps using the hierarchical clustering distance method with the function of heatmap, SOM (Self-organization mapping) and k-means using the silhouette coefficient to adapt the optimal classification with default parameters in R. Gene Ontology (GO) enrichment analysis of differentially expressed genes using the **GO**rilla (Gene **O**ntology en**R**ichment ana**L**ysis and vizua**L**iz**A**tion) tool (11, 12). GO terms with corrected *p*-values less than 0.05 were considered significantly enriched by differential expressed genes.

### Statistics

Unless stated otherwise all data shown are represented as mean ± standard error mean (SEM) of at least 3 biologically independent experiments. A *p*-value of ≤0.05 in an unpaired two-sided *t*-test or ANOVA test indicated a statistically significant difference. Kaplan-Meier estimates were calculated using the GraphPad Prism Software package. For Kaplan-Meier estimates a *p*-value of 0.05 in a log-rank test indicated a statistically significant difference.

## Results

### Quetiapine prevents radiation-induced phenotype conversion in vitro

We had previously reported that radiation treatment induced a phenotype conversion of glioma cells into glioma-initiating cells through re-expression of Yamanaka factors. Furthermore, that treatment with the first-generation dopamine receptor antagonist trifluoperazine prevented this phenotype conversion and prolonged survival in a syngeneic GL261 model of GBM as well as in patient-derived orthotopic xenografts (7).

Given the unfavorable side effect profile of trifluoperazine we screened a panel of additional 32 dopamine receptor antagonist for their ability to interfere with the process of radiation-induced phenotype conversion. Compared to trifluoperazine, the second-generation dopamine receptor antagonist quetiapine (QTP), known for a milder side effect profile (13), showed higher efficacy against radiation-induced phenotype conversion than trifluoperazine in this screening assay (**Figure 1A**).

**Figure 1.**
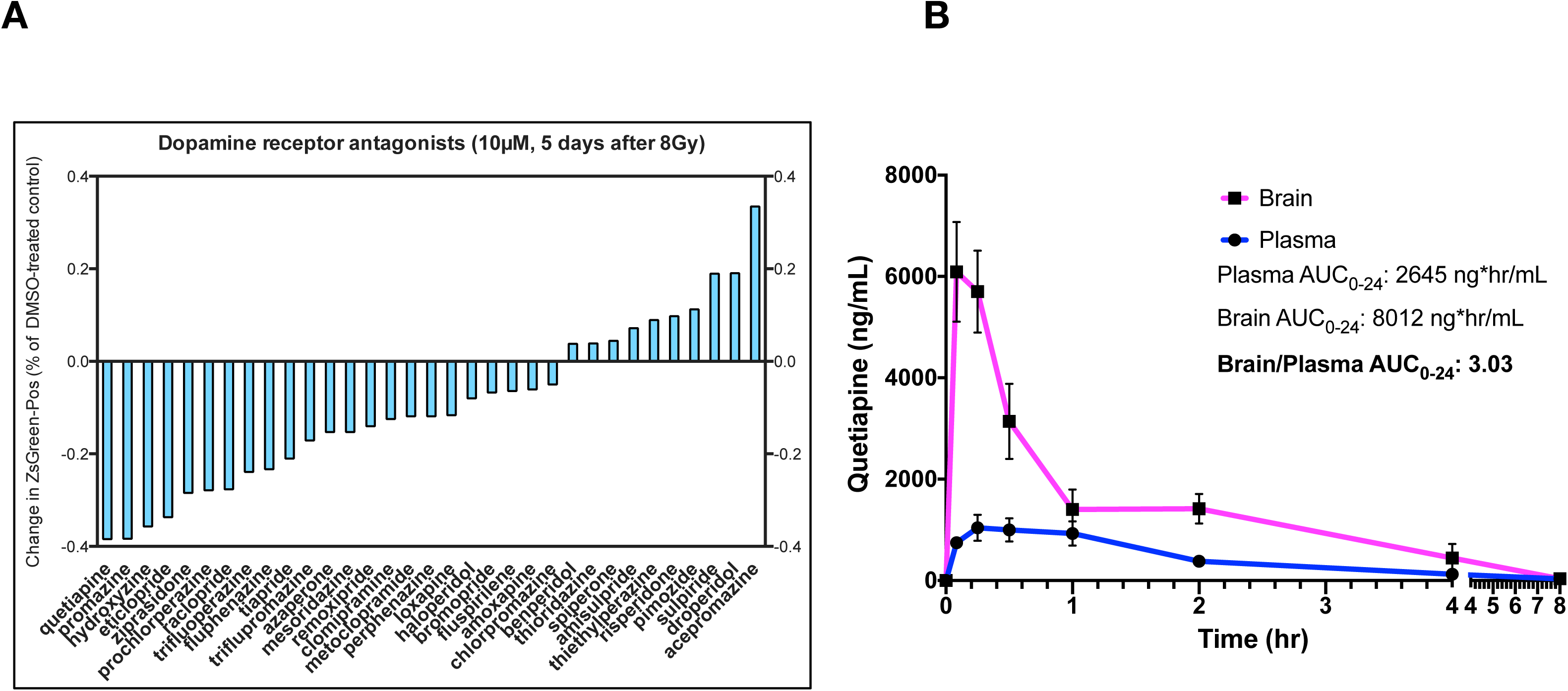
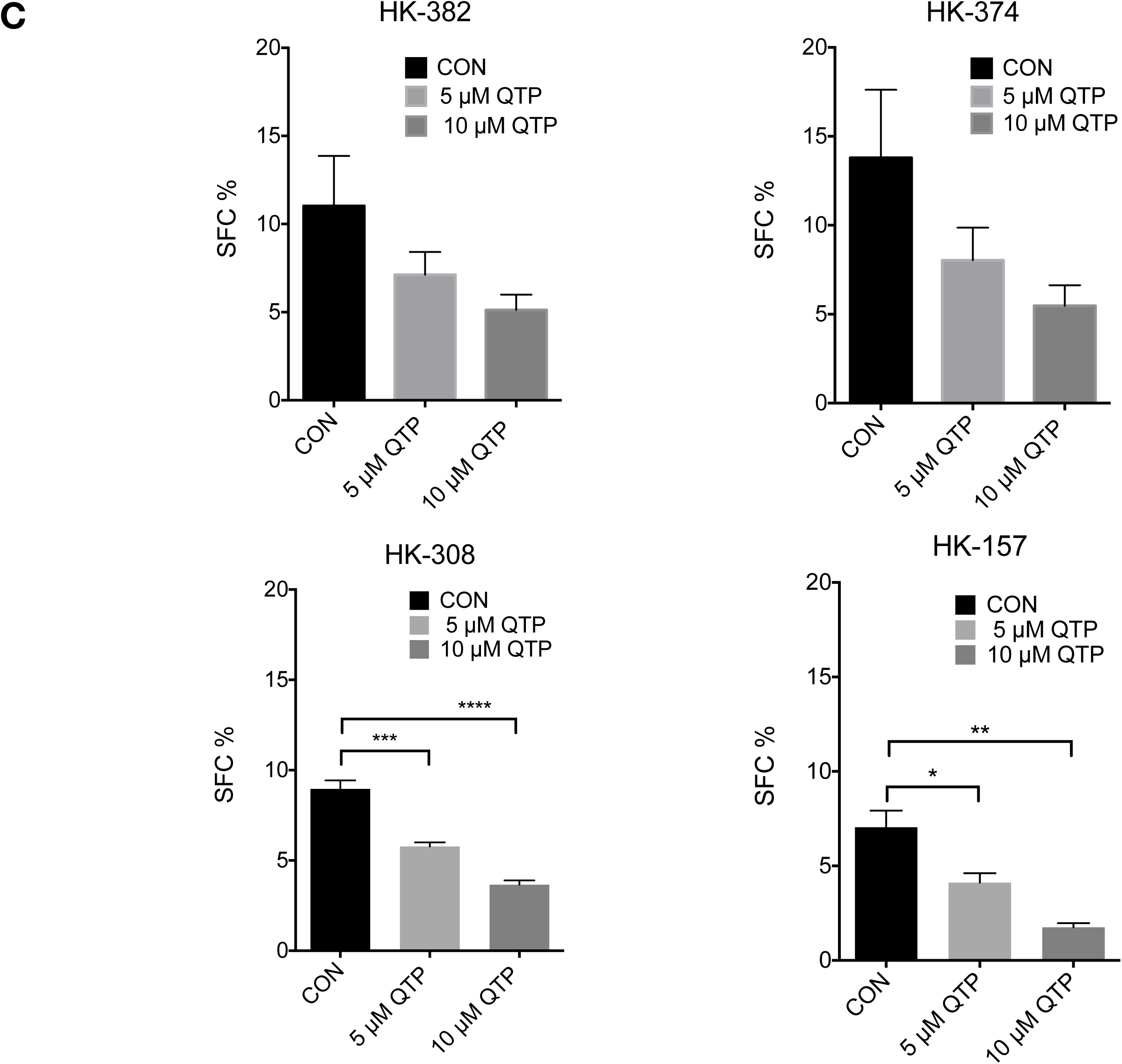
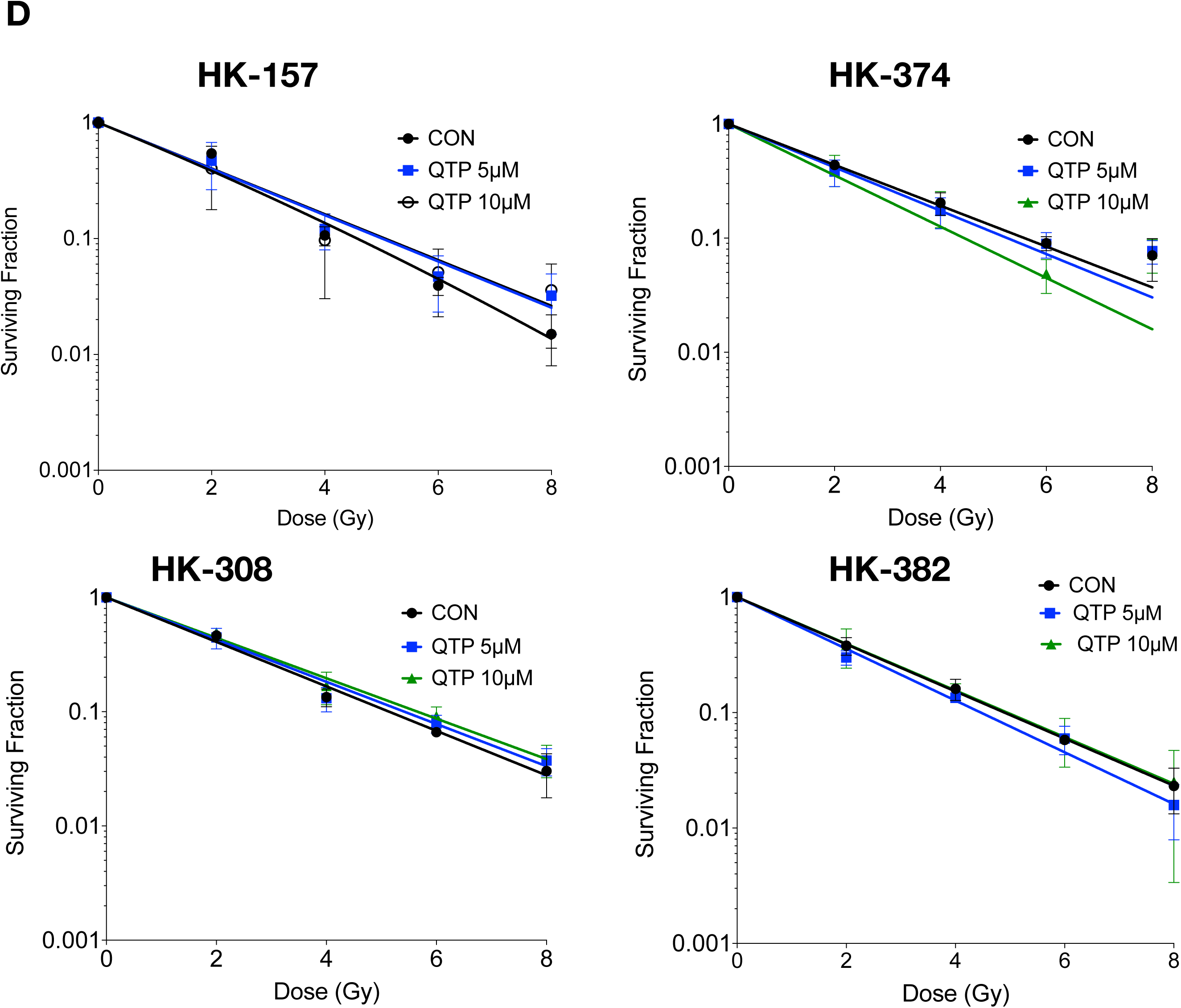
Quetiapine reduces self-renewal in patient-derived glioblastoma lines. **(A)** Inhibition of radiation-induced phenotype conversion by a panel of dopamine receptor antagonists. (**B**) Brain and plasma levels of quetiapine in mice after a single injection (30 mg/kg, i.p.). (**C**) Sphere-forming capacity of patient-derived glioblastoma lines in the presence or absence of quetiapine. (**D**) Surviving fraction of spheres treated with radiation in the presence or absence of quetiapine. All experiments in this figure have been performed with at least 3 independent biological repeats. (Unpaired t-test. * *p*-value < 0.05, ** *p*-value < 0.01, *** *p*-value < 0.001, **** *p*-value < 0.0001).

Even though QTP is a psychotropic drug we next ensured that QTP would penetrate into the CNS at relevant levels. Animals were treated with the mouse equivalent of 25% of the human MTD i.p. (30 mg/kg). Blood and brains were harvested at various time points after injection and subjected to mass-spectrometry. QTP rapidly crossed the blood brain barrier with a threefold higher AUC_0-24_ compared to plasma levels (**Figure 1B**).

In order to test the effects of QTP on the self-renewal of patient-derived glioblastoma specimens we treated glioma spheres of the HK-382, HK-374, HK-157, or HK-308 lines with five daily doses of QTP at 0, 5, or 10 *µ*M. QTP caused a dose dependent significant reduction in sphere-formation in all 4 specimens (**Figure 1C**). Next we combined single fractions of radiation with five daily doses of QTP to assess if QTP would act as a radiosensitizer in gliomaspheres. In none of the four patient-derived lines tested QTP sensitized glioma cells to radiation, suggesting that QTP treatment would not increase the toxicity of radiation (**Figure 1D**).

### Quetiapine prolongs survival in mouse models of glioblastoma

We next grafted GL261 cells into the striatum of C57BL/6 mice. Seven days after injection of the tumor cells we started treatment of the animals with escalating doses of QTP. Five daily doses of QTP per week for three weeks led to a significant gain in median survival (Control vs 30 mg/kg QTP: 23 vs 31 days, *p*=0.0181) but there was no apparent increase in survival correlated to the dose of QTP (**Figure 2A**). Therefore, we performed all subsequent experiments with QTP at 30mg/kg.

**Figure 2.**
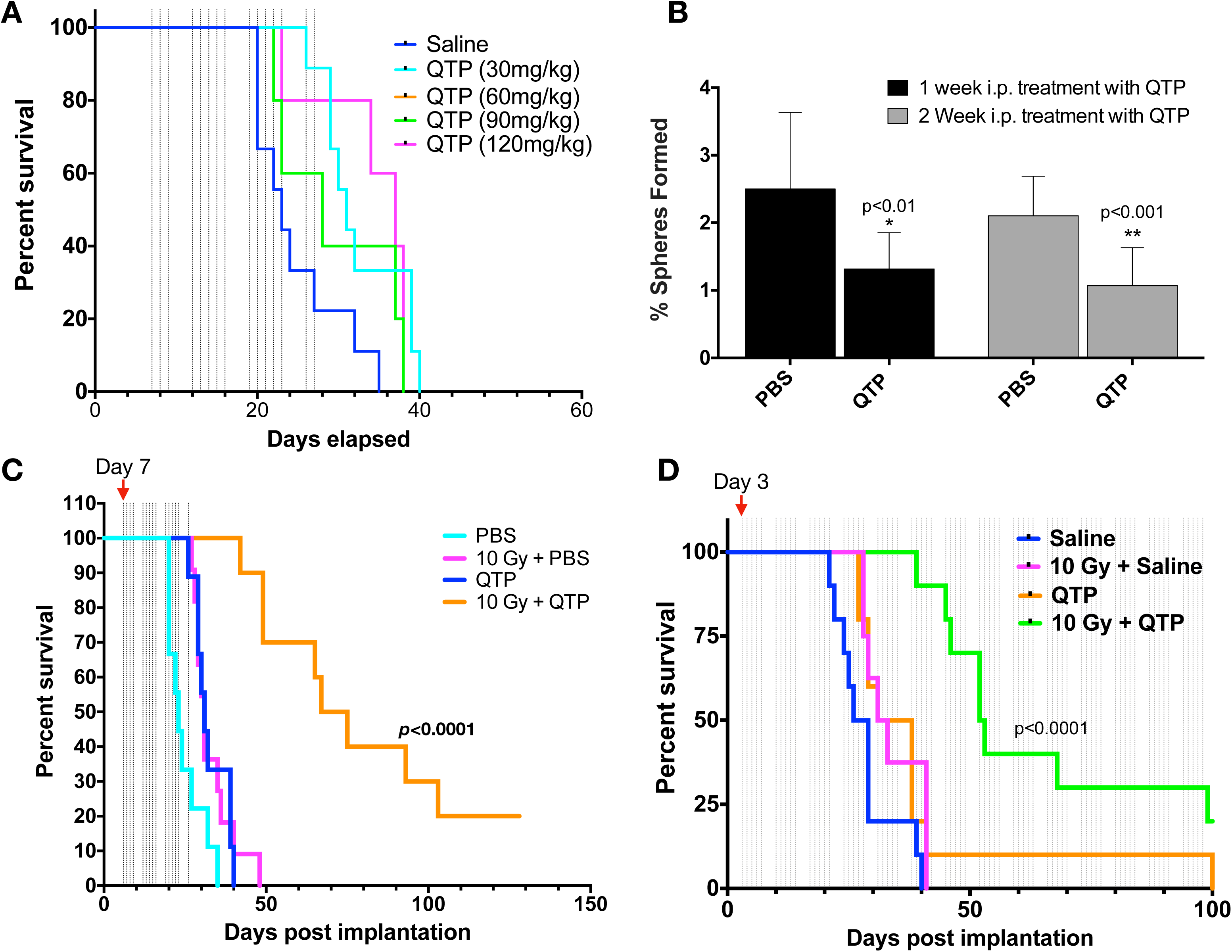
A combination of quetiapine and radiation prolongs survival in mouse models of glioblastoma. **(A)** Dose escalation of quetiapine in GL261 tumor-bearing C57Bl/6 mice. (**B**) Treatment of GL261 tumor-bearing C57Bl/6 mice with quetiapine reduces the sphere-forming capacity of surviving tumor cells. This experiment has been performed with at least 2 independent biological repeats. (Unpaired t-test. * *p*-value < 0.05, ** *p*-value < 0.01) (**C/D**) A combination of radiation (10 Gy) and daily injections of quetiapine significantly prolongs survival in the GL261 glioma mouse model (**C**) and in mice carrying patient-derived orthotopic xenografts (**D**). Log-Rank (Mantel-Cox) test for saline vs. 10 Gy + QTP *p*-value < 0.0001.

Treatment of mice bearing GL261-StrawberryRed tumors with 5 daily injections of QTP for one or two weeks significantly reduced the sphere-forming capacity of the surviving tumors cells, thus indicating efficacy of QTP against glioma-initiating cells *in vivo* (**Figure 2B**),

A single radiation dose of 10 Gy seven days after implantation of GL261 cells into the striatum of C57BL/6 mice led to a small increase in survival (Control versus 10 Gy: 23 vs 31 days, *p*=0.0126), comparable to the effect of QTP treatment (Control versus QTP: 23 vs 31 days, *p*=0.0181). However, irradiation with 10 Gy rendered the tumors vulnerable to five daily injections of QTP per week, and this treatment led to a significant increase in median survival from 23 to 71 days (*p*<0.0001) (**Figure 2C**).

Next we verified these findings using patient-derived orthotopic xenografts. Three days after implantation of HK-374 glioblastoma cells into the striatum of NSG mice, animals were treated with a single dose of 0 or 10 Gy followed by daily injection of QTP or saline (5 days on – 2 day off schedule). Although NSG mice are deficient in DNA-repair via non-homologous end-joining, normal brain tissue toxicity from radiation does not differ from that in immune-competent mice (14). Like in mice bearing GL261 tumors, QTP treatment or irradiation alone led to small increases in median survival (Control vs QTP: 27.5 vs 34.5 days, n.s.; Control vs 10 Gy: 27.5 vs 32 days, *p*=0.0451). However, the combination of a single radiation dose of 10 Gy followed by treatment with QTP increased the median survival from 27.5 to 52.5 days (log-rank test, *p*<0.0001) (**Figure 2 D**).

### Dopamine receptor inhibition in combination with radiation induces the expression of genes involved in cholesterol biosynthesis

While the combined treatment with radiation and QTP significantly improved the median survival of the animals, almost all animals eventually succumbed to the implanted tumor, thus suggesting that the tumor cells initiated a response to the treatment that led to resistance to QTP. To study this possibility in more detail we next performed RNA-Sequencing on HK-374 cells at 48 hours after irradiation and/or QTP treatment to access changes in the expression of genes in the response to combined treatment. When compared to samples receiving only a single radiation of 10 Gy, the combined treatment with radiation and QTP led to the differential up-regulation of 587 and the downregulation of 457 genes (**Figure 3A).** The combination of QTP and radiation led to the differential expression of 146 unique genes (**Figure 3B**) and irradiated samples showed a distinct genes expression profile (**Figure 3C**). The top 10 up- and downregulated genes are presented as a heatmap **(Figure 3D)** and were validated using qRT-PCR **(Figure 3E).** Gene Ontology enrichment analysis revealed overlap of the differentially down-regulated genes in the samples treated with radiation and QTP with gene sets involved in cell cycle checkpoint and cell cycle progression, chromatid segregation, G1/S transition of the cell cycle, RNA processing and DNA damage (**Figure 3F**).

**Figure 3.**
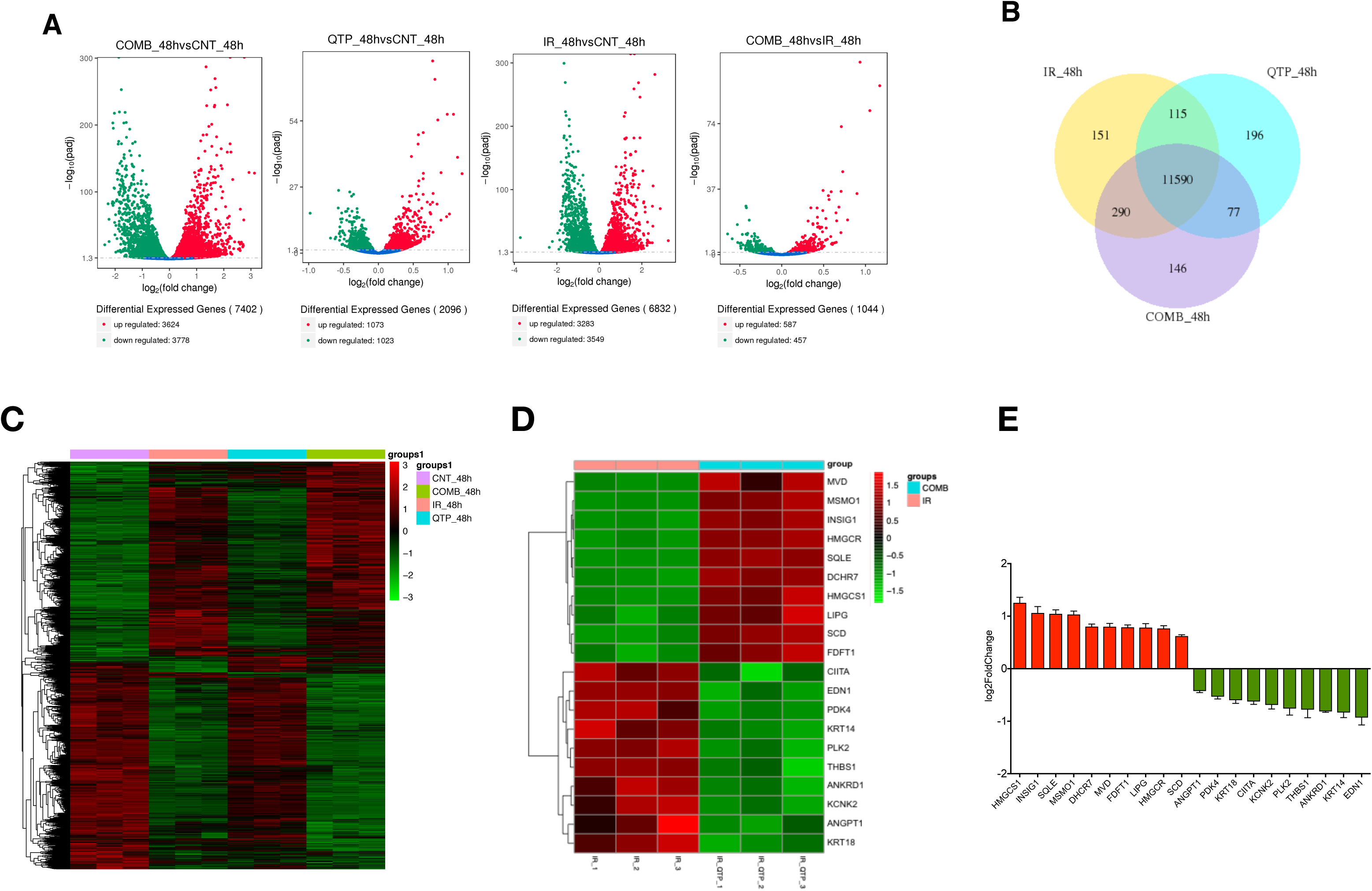
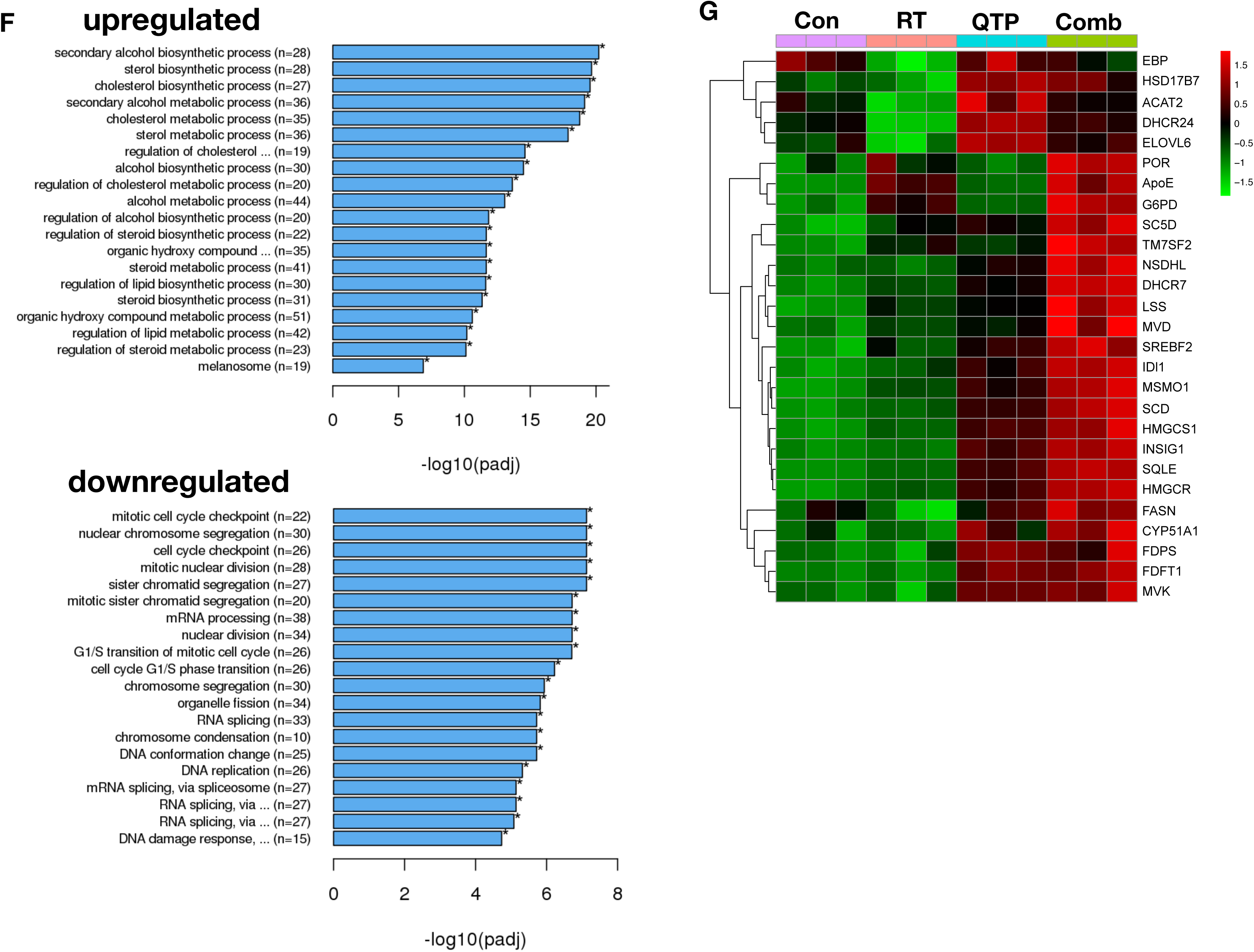
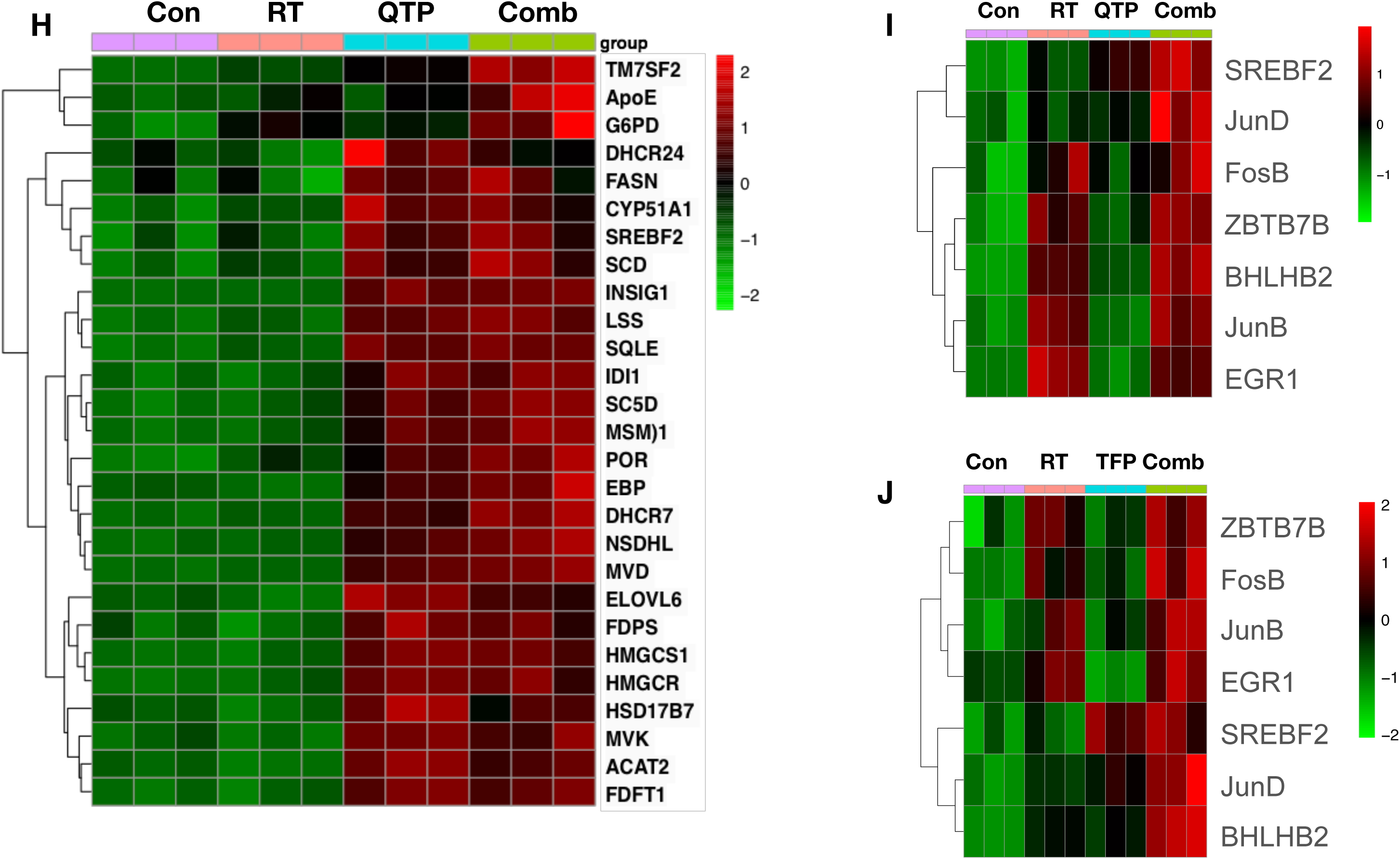
RNA-Seq analysis of glioma cells treated with QTP and radiation. (**A**) Volcano diagrams of differentially expressed genes in HK-374 cells, 48 hours after treatment with radiation (4 Gy), QTP or QTP and radiation compared to untreated control cells (**B**) Venn diagrams of differentially expressed genes in HK-374 cells, 48 hours after treatment with radiation (yellow), QTP (blue) or QTP and radiation (violet). (**C**) Heatmap of differential expression genes in HK-374 cells, 48 hours after treatment with radiation, QTP or QTP and radiation. (**D**) Top-ten up- and downregulated genes and their validation by qRT-PCR (**E**). Top-twenty Gene Ontology gene set overlapping with genes differentially up- or downregulated in HK-374 cells, 48 hours after treatment with radiation and quetiapine (**F**). Genes involved in the biosynthesis of cholesterol are upregulated in HK-374 cells, 48 hours after treatment with radiation (4 Gy) and quetiapine (**G**) or radiation and trifluoperazine (**H**). Genes of the first-level regulator network of SREBF2, the master regulator of cholesterol biosynthesis are upregulated in HK-374 cells, 48 hours after treatment with radiation (4 Gy) and quetiapine (**I**) or radiation and trifluoperazine (**J**).

The most prominent genes set that overlapped with genes differentially upregulated after combined treatment were involved in cholesterol, sterol and lipid biosynthesis (**Figure 3F**). The majority of genes involved in the cholesterol biosynthesis pathway were upregulated by QTP treatment, but this effect was enhanced when QTP treatment was combined with radiation (**Figure 3G**). Analysis of the same set of genes in a second RNA-Seq data set of HK-374 cells treated with radiation, TFP or a combination of TFP and radiation showed a similar upregulation of genes involved in cholesterol biosynthesis after combined treatment and trifluoperazine treatment alone (**Figure 3H**).

Most of the genes in this pathway are under the control of the transcription factor SREBF2 (sterol regulatory element binding transcription factor 2) (15) and our analysis found SREBF2 as well as several genes of SREBF2’s first level regulatory network upregulated in cells treated with QTP (**Figure 3I**) or Trifluoperazine and radiation (**Figure 3J**).

### Inhibition of cholesterol biosynthesis increases the efficacy of quetiapine and radiation against glioblastoma

Results from our RNA-Seq study led us to hypothesize that the upregulation of cholesterol biosynthesis is part of a defense mechanism of GBM cells against radiation combined with dopamine receptor inhibition. We therefore tested in treatment with QTP and radiation would render GBM cells vulnerable to treatment with the 3-Hydroxy-3-methylglutaryl-CoA reductase inhibitor Atorvastatin (ATR), a well-established and FDA-approved statin. Using the same four patient-derived GBM specimens in sphere-formation assays we reduced the very effective 5 daily treatments with QTP to only one treatment and combined it with ATR at 0.1, 0.25, 0.5 or 1 *µ*M concentrations. While a single dose of QTP had limited inhibitory effect on the self-renewal of gliomaspheres, the addition of ATR to QTP treatment led to a significant, dose-dependent reduction in sphere-formation (**Figure 4A**). When combined with radiation, the addition of ATR did not radiosensitize gliomaspheres in the presence or absence of QTP (**Figure 4B**). *In vivo*, daily treatment with QTP and ATR (5 days on – 2 days off schedule) after a single dose of 10 Gy (**Figure 4C**) significantly increased the survival of C57BL/6 mice implanted with GL261 cells with 90% of the animals surviving 157 days (*p*<0.0001) (**Figure 4D**). In NSG mice bearing HK-374 PDOXs the same combination treatment increased the median survival to 171 days (*p*<0.0001) (**Figure 4E**).

**Figure 4.**
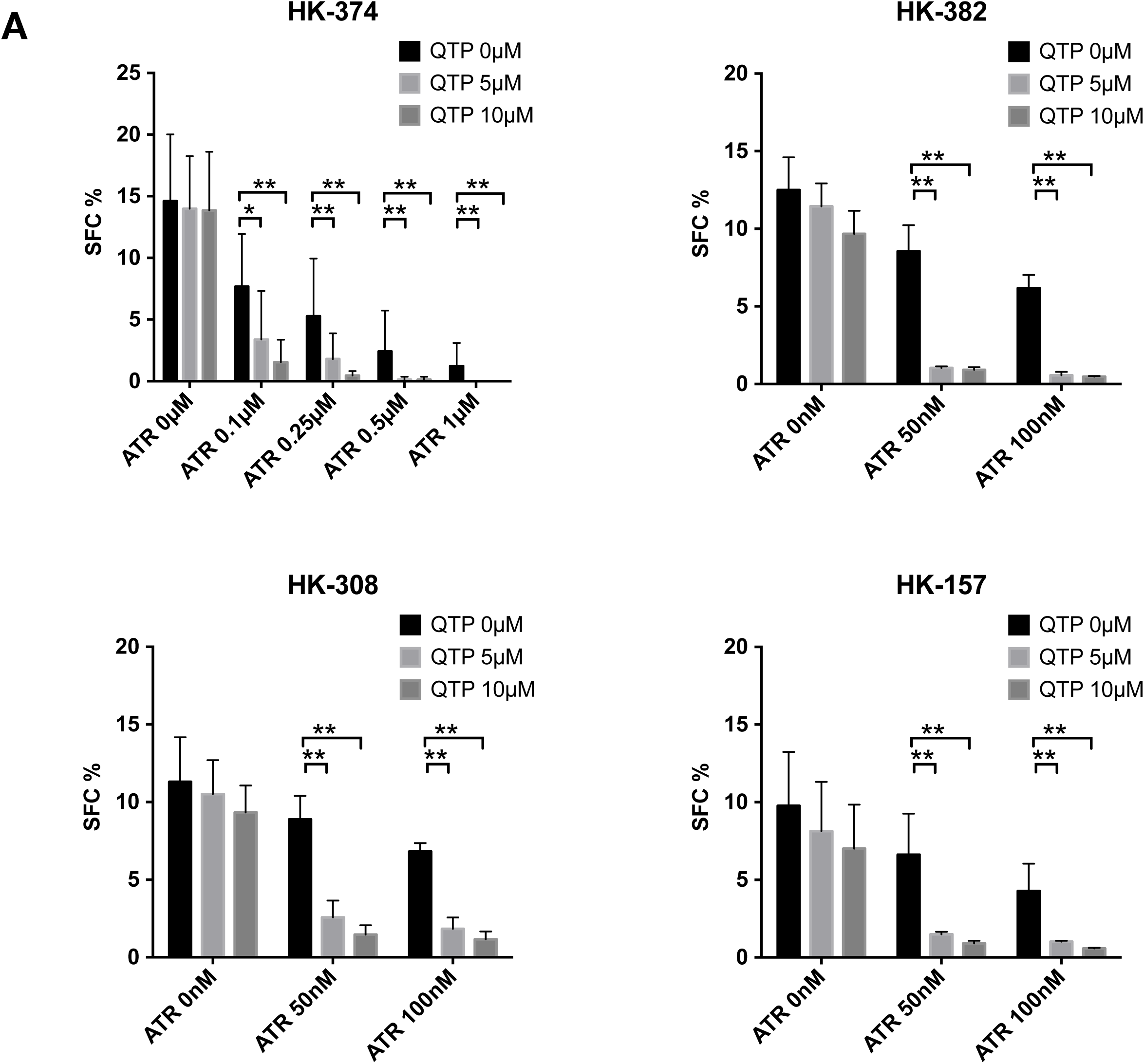
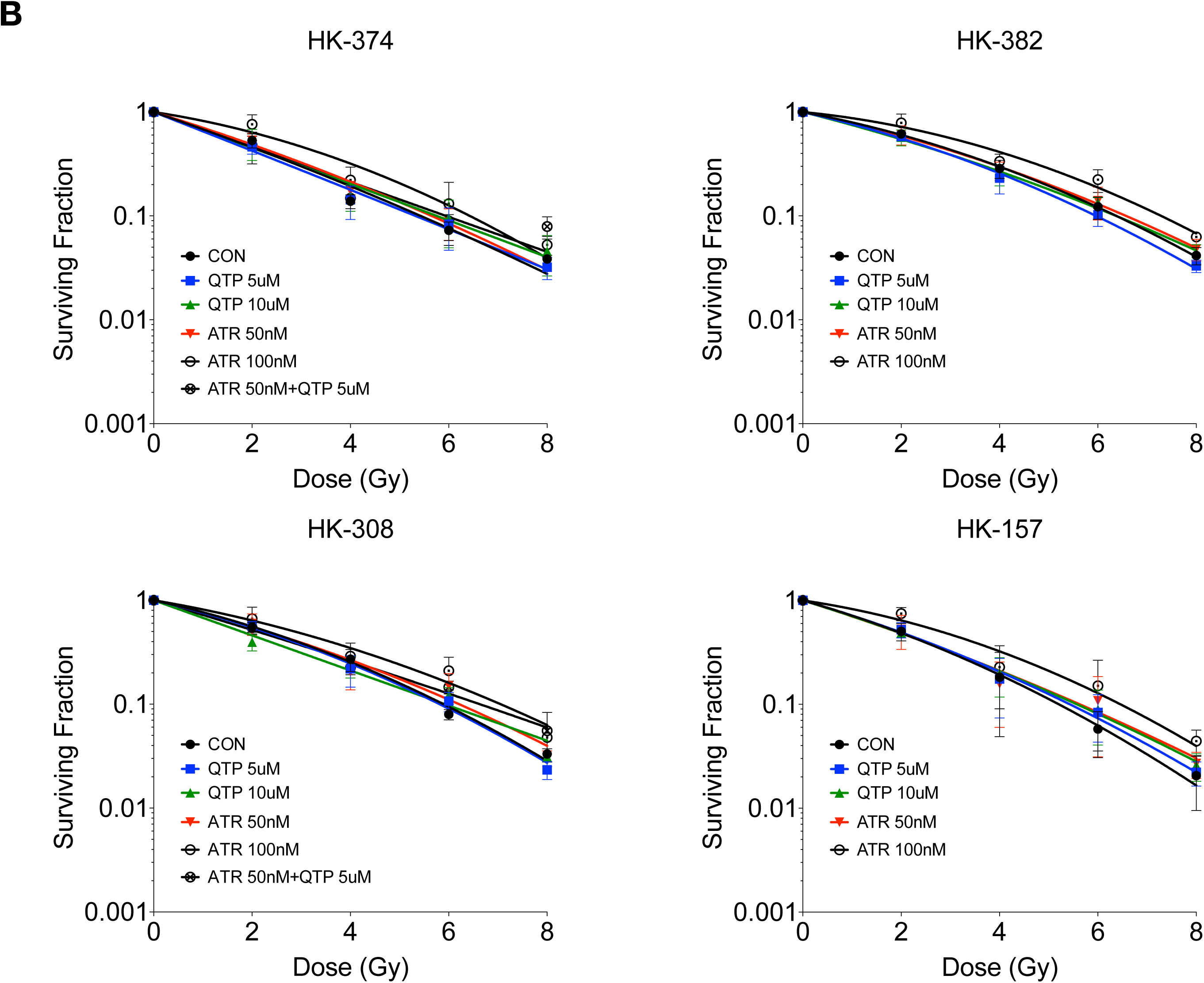
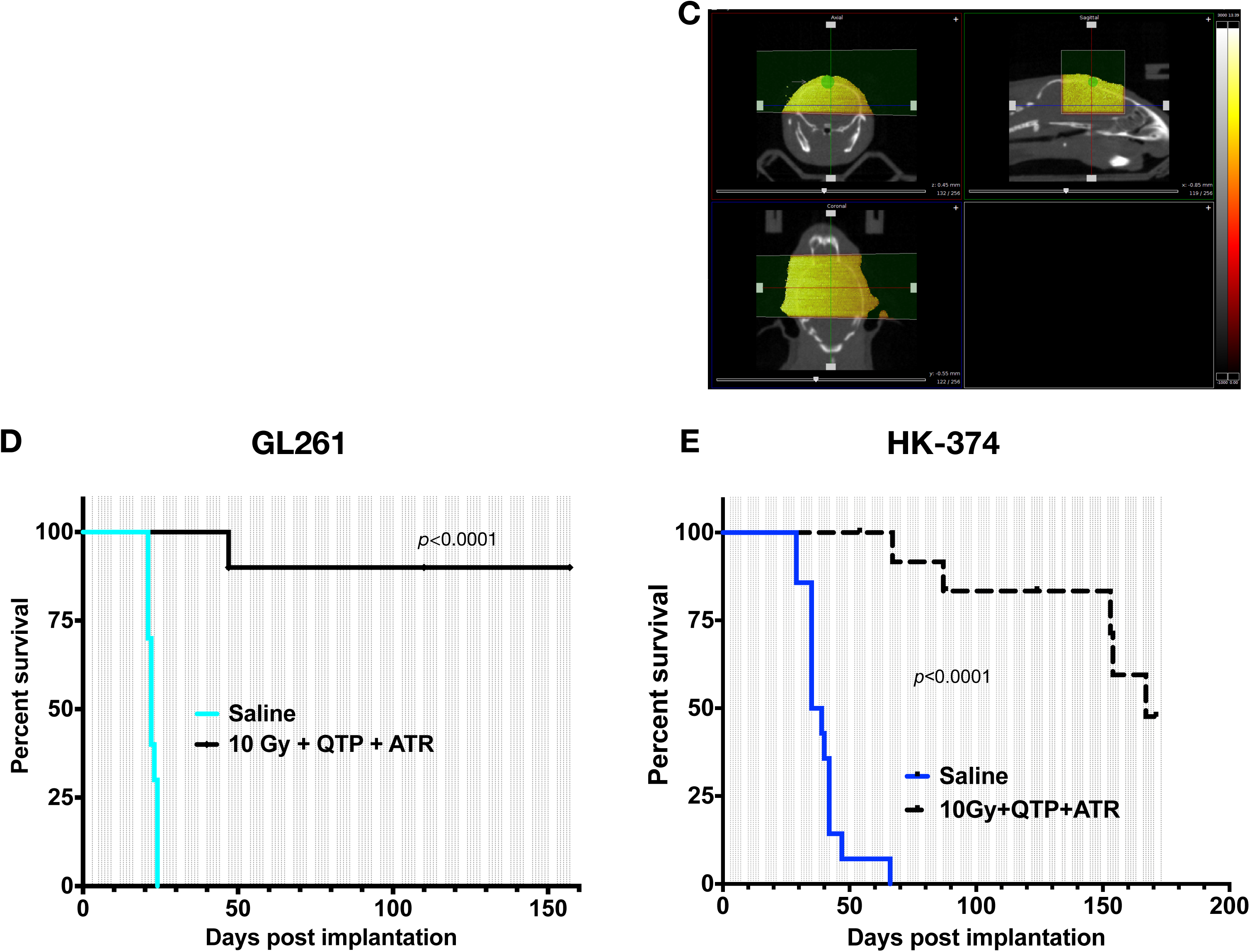
A combination of quetiapine and radiation with atorvastatin improves survival in mouse models of glioblastoma. (**A**) Quetiapine and atorvastatin synergistically decrease sphere-formation in patient-derived glioblastoma lines. All experiments in this figure have been performed with at least 3 independent biological repeats. (Unpaired two-way ANOVA. * *p*-value < 0.05, ** *p*-value < 0.01, *** *p*-value < 0.001, **** *p*-value < 0.0001). (**B**) Quetiapine, atorvastatin or a combination of both does not alter the radiation sensitivity of patient-derived glioblastoma lines. (**C**) Treatment plan for patient-derived orthotopic xenografts using an image-guided small animal irradiator. (**D**) A combination of quetiapine and atorvastatin (both 30 mg/kg) with a single dose of radiation (10 Gy) prolongs survival in the GL261 glioma mouse model (D) and HK-374 patient-derived orthotopic xenografts (**E**). Log-Rank (Mantel-Cox) test for saline vs. 10 Gy + QTP + ATR *p*-value < 0.0001.

## Discussion

One of the main pillars in the standard-of-care for patients suffering from glioblastoma is radiotherapy and its addition to the treatment regimens is one of the few modalities that robustly prolong the survival of the patients over surgery alone (16). Out of many attempts that added conventional chemotherapy or targeted therapies to the standard-of-care so far only temozolomide met the threshold to be included into the treatment regimen (17). However, despite decades of effort the overall survival of patients with glioblastoma is still unacceptably low, thus indicating an unmet need for novel treatment options against glioblastoma.

Recent reports in the literature have suggested an anti-tumor activity of dopamine receptor type 2 antagonists against glioblastoma (18-21) and dopamine receptors have been found to be expressed in various tumor types including glioblastoma (22). However, the anti-tumor efficacy of dopamine receptor antagonists as single agents was found to be limited in preclinical (23) and clinical studies (24).

We had recently reported that a combination of a single high dose of radiation and continuous application of the first-generation dopamine receptor antagonist trifluoperazine improved survival in mouse models of glioblastoma (7). Here we report that QTP, a second-generation dopamine receptor antagonist with a more favorable side effect profile than trifluoperazine, when combined with radiation also shows efficacy against glioblastoma-initiating cells *in vitro* and prolongs survival in mouse models of glioblastoma without affecting the radiation sensitivity of glioma cells. While the effect on survival was highly significant and meaningful, most animals ultimately showed tumor progression and our search for a possible resistance mechanism revealed upregulation of key components of the cholesterol biosynthesis pathway in the response of glioblastoma cells to the combined treatment with dopamine receptor antagonists and radiation. A previous report had shown dependence of the survival of glioblastoma cells on cholesterol (25) but a potential beneficial impact of statin use in glioblastoma patients is currently discussed controversially (26-29). We report here that QTP treatment but not irradiation alone led to upregulation of the key components of the cholesterol biosynthesis. This effect was amplified when QTP and radiation were combined. And while inhibition of cholesterol biosynthesis using ATR by itself had some effect on glioblastoma cells, the combined treatment with radiation and QTP rendered the cells and tumors particularly sensitive to the addition of ATR.

In summary we conclude that dopamine receptor antagonists are readily-available, FDA-approved drugs with well-known toxicity profiles that enhance the efficacy of radiotherapy in glioblastoma, especially in combination with the statin atorvastatin, a treatment combination that can be easily tested in future clinical trials.

## Acknowledgements

FP was supported by grants from the *National Cancer Institute* (R01CA137110, R01CA161294). DN, PLN, TC, LL, HK, and FP were supported by a grant from the *National Cancer Institute* (P50CA211015).

---

**Supplementary Table 1.**
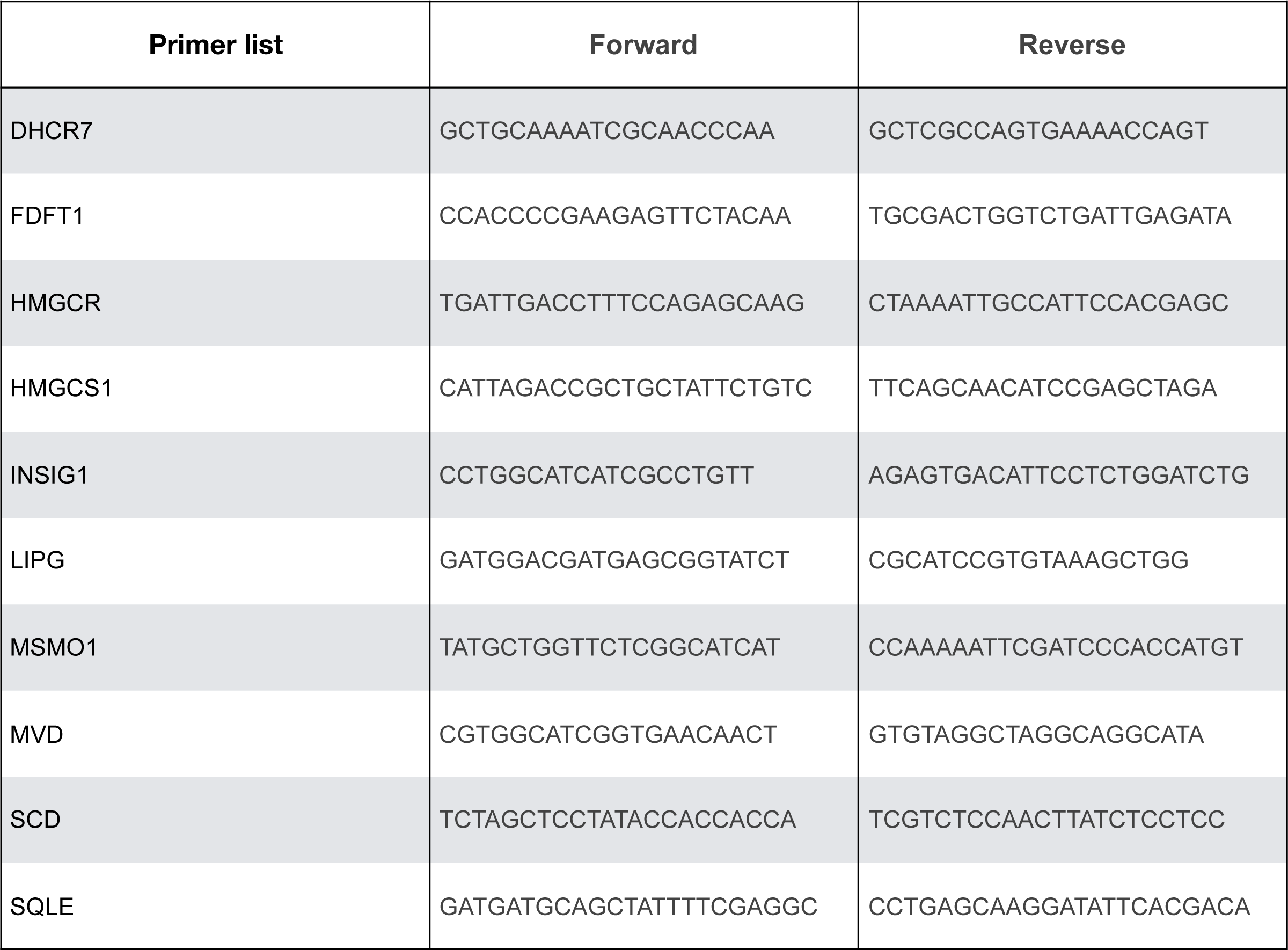

